# A putative *cis*-regulatory element near *Shox2* in the jerboa genome (*Jaculus jaculus*) is not sufficient to alter mouse limb bone lengths

**DOI:** 10.64898/2026.09.17.752424

**Authors:** Galilea Guerrero, Alexander Y. Liu, Alexander J. Weitzel, Erica Gacasan, Michael Hiller, Robert Sah, Kimberly L. Cooper

## Abstract

Vertebrate limb bones grow at different rates within and between species to achieve a striking variety of proportions and functions. However, most coding mutations that alter bone growth in mice and humans affect the entire skeleton, pointing to *cis-*regulatory modularity as the primary mechanism to diversify proportion. In a search for causative loci, we previously identified 1,755 disproportionately differentially expressed genes associated with the extreme hindlimb elongation in bipedal jerboas, *Jaculus jaculus*, compared to laboratory mice. We then took a focused approach to identify putative *cis*-regulatory elements near select candidate genes in a highly fragmented draft *Jaculus jaculus* genome assembly. We identified an ATAC-seq peak near *Shox2* that was reproducible in jerboa metatarsals and not in radius/ulna. *Shox2* gained a novel expression domain in jerboa metatarsals and is both necessary and sufficient to modulate mouse limb bone length. This 139 bp putative enhancer in the jerboa genome centers on a ∼1 kb deletion of sequence found broadly throughout placental mammals. Here, we show that replicating the deletion and replacement with the 139 bp jerboa sequence in mice causes no detected phenotype. A genome-wide analysis of our ATAC-seq data using a high-quality *Jaculus jaculus* genome assembly replicated the peak called at this location but also identified 7,445 peaks that are similarly reproducible in metatarsals and not in radius/ulna. Certainly not all of these peaks are causative, but it is likely that many small-effect mutations diversified proportion. It is therefore crucial to apply unbiased filters to genome-wide analyses and to not assume that the few historically favored genes will have the largest effect.

## Introduction

Vertebrate animals navigate their environment using a striking diversity of limb shapes and sizes. The forelimb bones of a mole, for example, are short and robust to act as shovels tunneling through dirt while the foot bones of running mammals are elongated for greater stride length. However, early embryonic limb rudiments are far more similar to one another, and individual bones grow at different rates and durations to establish the diversity of adult skeletal proportions (Fröbisch & Shubin, 2011; Hockman et al., 2009; Keenan & Beck, 2016; Martin, 1990; Montero et al., 2017).

Coding sequence mutations in genes that regulate bone development frequently cause dwarfism throughout the skeleton (Liu et al., 1993; Watanabe et al., 1994; Zhou et al., 1997) suggesting a common ‘toolkit’ regulates bone growth. Pleiotropic gene function therefore indicates that development and diversification of limb proportion results from millions of years of enhancer evolution. However, *cis-*regulatory mechanisms that locally ‘tune’ skeletal growth to achieve adult proportion and to diversify its evolution are currently not well understood. These are likely modular in the sense that multiple enhancers control expression of a gene throughout the skeleton by differential usage of individual enhancers, or of transcription factor binding sites within an enhancer, in different locations.

To uncover the genetic basis of modular bone growth, we use the bipedal lesser Egyptian jerboa (*Jaculus jaculus*), a desert-adapted jumping rodent with strikingly elongated hindlimbs and disproportionately long metatarsals. Among the closest relatives of laboratory mice, for which we understand the most about mechanisms of limb growth, jerboas have the most extreme limb morphology. Further, jerboas retained a similar ‘mouse-like’ forelimb, which allows us to identify and exclude genetic divergence that is unrelated to the evolution of hindlimb skeletal proportion (Saxena et al., 2022).

We first sought to identify genes that are associated with disproportionate growth by performing interspecies transcriptomic analyses comparing jerboa and mouse metatarsals of the hindfoot and radius/ulna of the forearm. We identified 1,755 genes, ∼10% of orthologs, that are disproportionately differentially expressed in jerboa metatarsals compared with radius/ulna (Saxena et al., 2022). These genes are likely controlled by modular enhancers in the skeleton, though expression differences between species may be due to *cis-*regulatory sequence divergence or differences in expression of upstream transcription factors.

To first identify candidate *cis*-regulatory sequences, we performed ATAC-seq analysis using the *Jaculus jaculus* draft genome (JacJac1.0) consisting of 10,899 scaffolds assembled from Illumina Hi-Seq reads. The incompleteness and fragmentation of this assembly limited our ability to assign all peaks to nearest genes. We therefore identified all significantly accessible and reproducible peaks (MACS2 and IDR) in jerboa metatarsal and radius/ulna and searched for metatarsal ‘unique’ peaks within a genomic window spanning 500 kb from the transcriptional start sites of a few key differentially expressed candidate genes (Saxena et al., 2022).

We identified an ATAC-seq peak near *Short stature homeobox 2* (*Shox2*) that was reproducible in jerboa metatarsals but not radius/ulna. RNA-Scope *in situ* hybridization confirmed that jerboas gained a domain of *Shox2* expression in the elongated metatarsal growth plates, where its expression is absent in mice (Clement-Jones et al., 2000; Cobb et al., 2006; Neufeld et al., 2014; Yu et al., 2007). Consistent with a hypothesis that *Shox2* contributes to jerboa metatarsal elongation, we also showed that misexpression of *Shox2* in mouse limbs is sufficient to promote distal limb elongation (Saxena et al., 2022).

The 139 bp ATAC-seq peak in the jerboa genome lies ∼285 kb upstream of the transcription start site of *Shox2* in an intron of the neighboring gene, *Rscr1*. *Rscr1* harbors a previously validated enhancer of *Shox2* in mice and humans (m741/hs741) (Osterwalder et al., 2018; Ye et al., 2016) and is itself not disproportionately differentially expressed in jerboa metatarsals. The sequence orthologous to the jerboa peak in the mouse genome was not accessible in either mouse growth cartilage (Saxena et al., 2022).

*Shox* genes are necessary for proximal limb bone elongation in humans and in mice. In humans, the *Short stature homeobox* (*SHOX*) gene controls growth of zeugopodal elements (forearm and lower leg), and mutations cause short stature (Clement-Jones et al., 2000). *SHOX* haploinsufficiency in patients with Turner Syndrome causes a slight reduction in the length of zeugopodal elements as well as cubitus valgus or “turned-out elbow”, a deformity characterized by the outward angling of the forearm relative to the body (Clement-Jones et al., 2000). Heterozygous *SHOX* mutations are associated with Léri–Weill dyschondrosteosis (LWD), characterized by shortening of zeugopodal elements and Madelung wrist deformities (Seki et al., 2014; Shears et al., 2002). Loss-of-function mutations in *SHOX* are associated with Langer Mesomelic Dysplasia (LMD), which causes the most drastic reduction in zeugopodal limb length and severely bowed limbs (Clement-Jones et al., 2000; Rao et al., 2001).

*Shox* has been lost from the mouse genome, but the *Shox2* paralog is retained with 99% amino acid sequence identity to human SHOX (Blaschke et al., 1998; Semina et al., 1998; Yu et al., 2007). *Shox2* is expressed more proximally in the limb in humans (Clement-Jones et al., 2000), chickens (Tiecke et al., 2006), and mice (Clement-Jones et al., 2000; Cobb et al., 2006; Neufeld et al., 2014; Yu et al., 2007). Consistent with its expression, the humerus and femur are shortened the most in *Shox2* mutant mice (Bobick & Cobb, 2012). The *Shox2* transcription factor binds to limb enhancers (Ye et al., 2016) and is required for chondrocyte differentiation and hypertrophy to promote long bone elongation (Bobick & Cobb, 2012; Yu et al., 2007). Together, these data suggested that the putative *cis*-regulatory sequence in the jerboa genome, preferentially accessible in metatarsals, could be responsible for gain of *Shox2* expression in jerboa metatarsals and might promote their disproportionate elongation.

Furthermore, multiple sequence alignment revealed that the 139 bp sequence in the jerboa ATAC-seq peak aligns to a 1228 bp sequence in mice (chr3:67266489-67267716). Although the underlying nucleotide sequence is not highly conserved, this greater length is conserved in an alignment of 40 placental mammals (Saxena et al., 2022). The *Jaculus jaculus* genome therefore has a ∼1 kb deletion at the site of this putative *cis*-regulatory element, which suggests evolution to a new function. Our analysis here further shows the deletion occurred prior to the last common ancestor of jerboas and jumping mice, which have moderately elongated hindlimbs to support facultative bipedalism.

To test the hypothesis that the jerboa 139 bp sequence is sufficient to promote *Shox2* expression and lengthen the distal hindlimb skeleton, we engineered mice that recapitulate the sequence found in jerboas at the orthologous location in the mouse genome. Specifically, we engineered the homozygous deletion of 1228 bp and replacement with the 139 bp jerboa sequence. However, we found no significant difference in expression of *Shox2* or bone lengths in the limbs of modified mice compared with wild type littermates.

Using a revised long read-based *J jaculus* assembly generated as part of the Vertebrate Genomes Project Phase I (Formenti et al., 2026), we re-analyzed the original ATAC-seq data genome-wide. Using the same criteria for significance and reproducibility in metatarsals but not radius/ulna, we again find the *Shox2-*associated peak and 7,445 additional loci. It is therefore difficult to determine if the *Shox2*-associated sequence does not contribute to skeletal proportion or if the single locus replacement is not sufficient. Ongoing work will apply functional and comparative genomics in an unbiased approach to refine the set of candidate loci. However, we expect that skeletal proportion behaves as a quantitative trait caused by many loci of small effect and largely in under-studied genes. Although it is currently not feasible to fully replicate such complex traits in mice, and it will be a challenge to select loci to model, thoughtful persistence will reveal smaller pieces of a larger puzzle.

## Results

We first expanded the comparative genomic analysis of the region containing the jerboa *Shox2*-associated peak to a multiple sequence alignment of 488 placental mammals. This alignment includes five bipedal jerboa species and the facultative bipedal meadow jumping mouse (*Zapus hudsonius*). *Z*. *hudsonius* shares the most recent common ancestor with all jerboas in the superfamily Dipodoidea, and it has hindlimbs that are intermediate in length between quadrupedal rodents and jerboas (Moore et al., 2015). The length of sequence at this locus is largely conserved across placental mammals, and the deletion associated with the jerboa peak is shared among all sequence d bipedal species of Dipodoidea (Figure 1 and Supplementary Figure S1).

**Figure 1.**
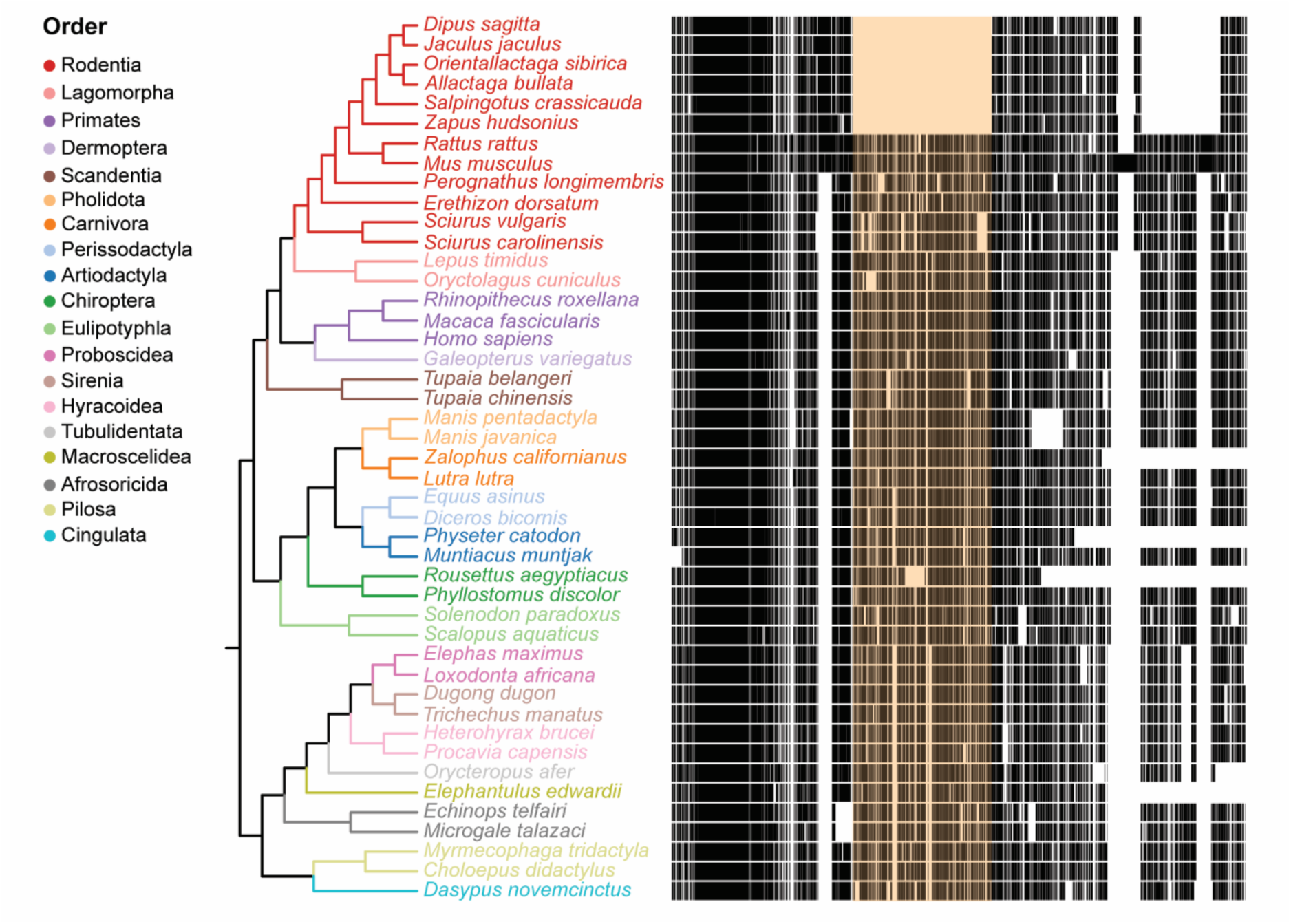
The ∼1 kb deletion is uniquely shared in the *Dipodoidea* superfamily. Alignment of the sequence flanking the identified ATAC-seq peak in a subset of 45 mammals reveals a 1089 bp deletion (highlighted in orange) that is shared among *Dipodoidea*, which includes jerboas and jumping mice, but not in other placental mammals. See the full alignment of 488 species in Supplemental Figure S1.

Due to compelling evidence that the 139 bp sequence under this jerboa ATAC-seq peak may harbor a *Shox2 cis*-regulatory element unique to these bipedal rodents, we recapitulated the genomic sequence configuration in mice. We used CRISPR to target sites flanking the 1228 bp orthologous region in the mouse genome and seamlessly replaced the deletion with the 139 bp jerboa sequence by homology directed repair using a single stranded oligodeoxynucleotide template with flanking mouse genomic homology and edited PAM sites to avoid re-cutting after insertion (Figure 2). This was a bespoke peak not identified from a larger dataset, a long series of numbers defines its genomic coordinates, and it is not a definitive regulator of *Shox2* expression. We therefore refer to these modified mice as ‘1228bp^ΔJ^’ mice. This indicates both the deletion and a 139 bp insertion of jerboa sequence at the peak locus. Heterozygous 1228bp^ΔJ/WT^ mice produced litters with expected Mendelian ratios (Supplemental Table S1), and heterozygotes and homozygotes have no obvious health issues and survive for a typical lifespan.

**Figure 2.**
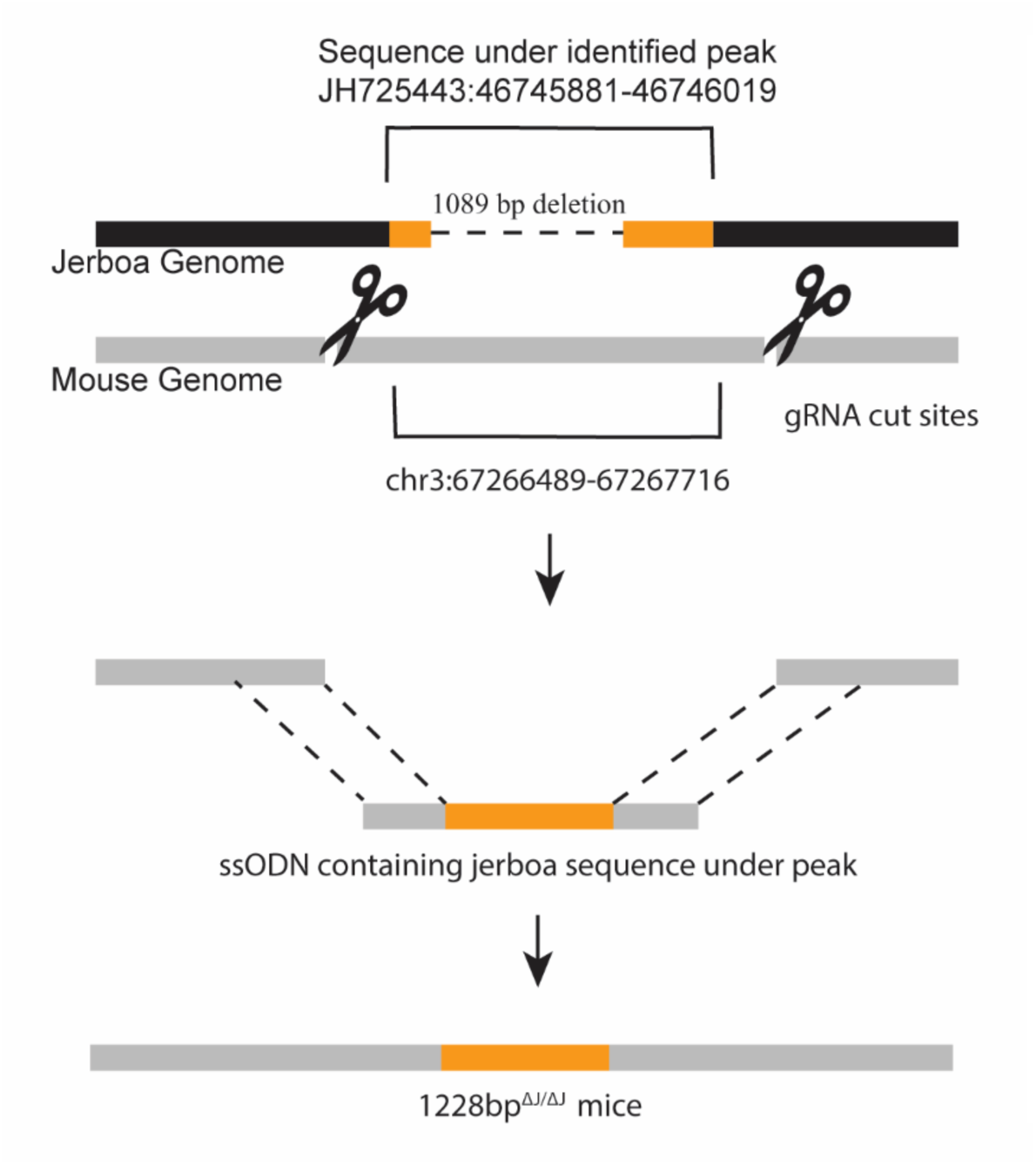
The sequence under the jerboa peak was recapitulated in 1228bp^ΔJ/ΔJ^ mice. Mice were modified to recapitulate the 139 bp sequence under a jerboa ATAC-seq peak identified ∼285 kb upstream of *Shox2* in an intron of *Rsrc1*. Two Cas9-mediated DSBs flanking the homologous region were introduced at chr3:67266338 and 67267772. The 139 bp jerboa peak sequence was inserted by homology-directed repair using an ssODN, which also recodes the PAM gRNA sequences, resulting in a 1228 bp deletion and replacement with the 139 bp jerboa ATAC-seq peak sequence in 1228bp^ΔJ/ΔJ^ mice.

To detect a disproportionate rate of limb bone elongation or disproportionate adult bone lengths that may be caused by the jerboa sequence replacement, we collected 1228bp^ΔJ/ΔJ^ homozygous mice and their wild type littermates (1228bp^WT/WT^) from birth (P0) to skeletal maturity (P42). Mice aged P14-P42 were scanned by micro-computed tomography (µCT). Due to the immature ossification state of neonatal limbs, µCT scans are unable to provide sufficient resolution to accurately measure bone lengths. We therefore stained P0 and P7 skeletons with alizarin red and alcian blue and measured their limb bones from digital images. For all specimens, we measured the length of skeletal elements in the hindlimb (femur, tibia, metatarsals), forelimb (humerus, radius, ulna, third metacarpal), and the naso-occipital length of the skull.

Limb bone lengths in 1228bp^ΔJ/ΔJ^ mice, absolute and normalized to skull length, are not significantly different from 1228bp^WT/WT^ controls at any age, and thus they follow the same growth trajectory throughout development (Figure 3, Supplemental Figures S2, S3). For the variance observed at the timepoint with largest difference between 1228bp^ΔJ/ΔJ^ and control, ∼4% in tibia at P42, we would need 22 mice per genotype per timepoint (220 mice) to achieve 90% power to determine statistical significance. The extremely divergent sequence associated with the jerboa metatarsal peak is therefore not sufficient to alter limb bone lengths or proportion to a reasonably measurable extent.

**Figure 3.**
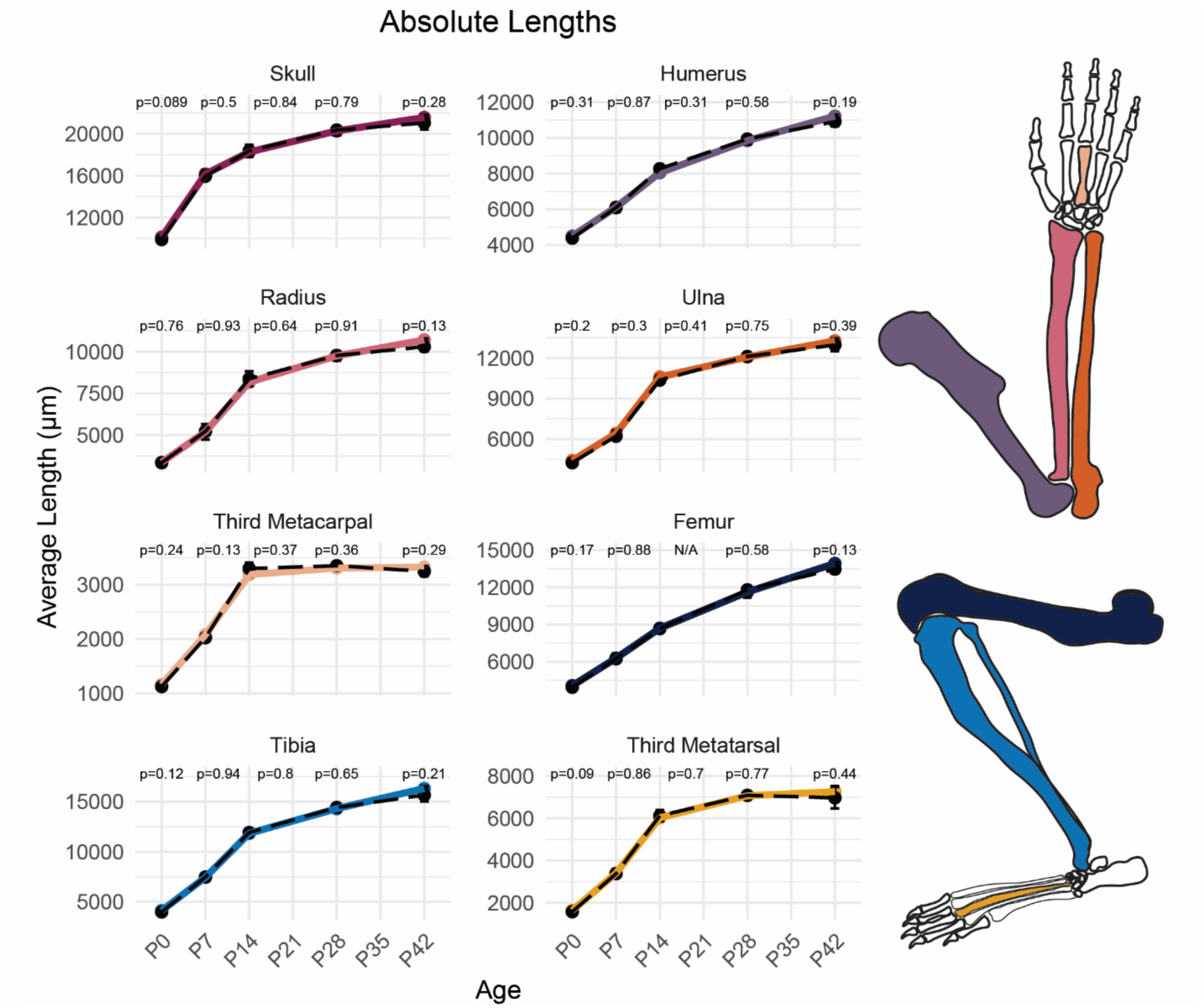
Modified mice have no detected limb skeletal phenotype. Lengths of the skull and limb skeletal elements were measured in modified 1228bp^ΔJ/ΔJ^ mice and wild type littermates at intervals from birth (P0) to adulthood (P42). Each measurement for one animal is the average length of the left and right limb skeletal elements. Wild type growth curves are represented by a black dashed line; 1228bp^ΔJ/ΔJ^ growth curves are presented as a solid color line. Statistical significance was determined by Student’s t-test. n≥ 3.

Although 1228bp^ΔJ/ΔJ^ mice did not exhibit a difference in limb length, the jerboa peak sequence might increase *Shox2* expression in mouse metatarsals below a level sufficient to increase bone length. *Shox2* has not been detected in mouse metatarsals (Cobb et al., 2006; Neufeld et al., 2014; Saxena et al., 2022), but we observed *Shox2* expression in the proliferative zone and perichondrium of jerboa metatarsals by *in situ* hybridization and by quantitative reverse transcriptase PCR (Saxena et al., 2022). The level of *Shox2* expression in jerboa metatarsals was comparable to expression in the radius/ulna. We also showed that experimental misexpression of *Shox2* in mouse limbs using a *Prx1*-driven conditional system, which increased *Shox2* expression 2-2.5 fold over wild type, was sufficient to lengthen the adult mouse metatarsal by 9% (Saxena et al., 2022).

We obtained radius/ulna and metatarsal growth plates from five-day old 1228bp^ΔJ/ΔJ^ and 1228bp^WT/WT^ mice for qPCR. We previously detected *Shox2* expression in jerboa metatarsals at this stage when jerboa metatarsal elongation has also accelerated relative to the mouse. However, there is no detectable *Shox2* expression difference in either the radius/ulna or metatarsals of 1228bp^ΔJ/ΔJ^ mice compared to wild type littermates (Figure 4). The sequence under the identified jerboa ATAC-seq peak is therefore not sufficient to significantly alter *Shox2* expression during early metatarsal elongation.

**Figure 4.**
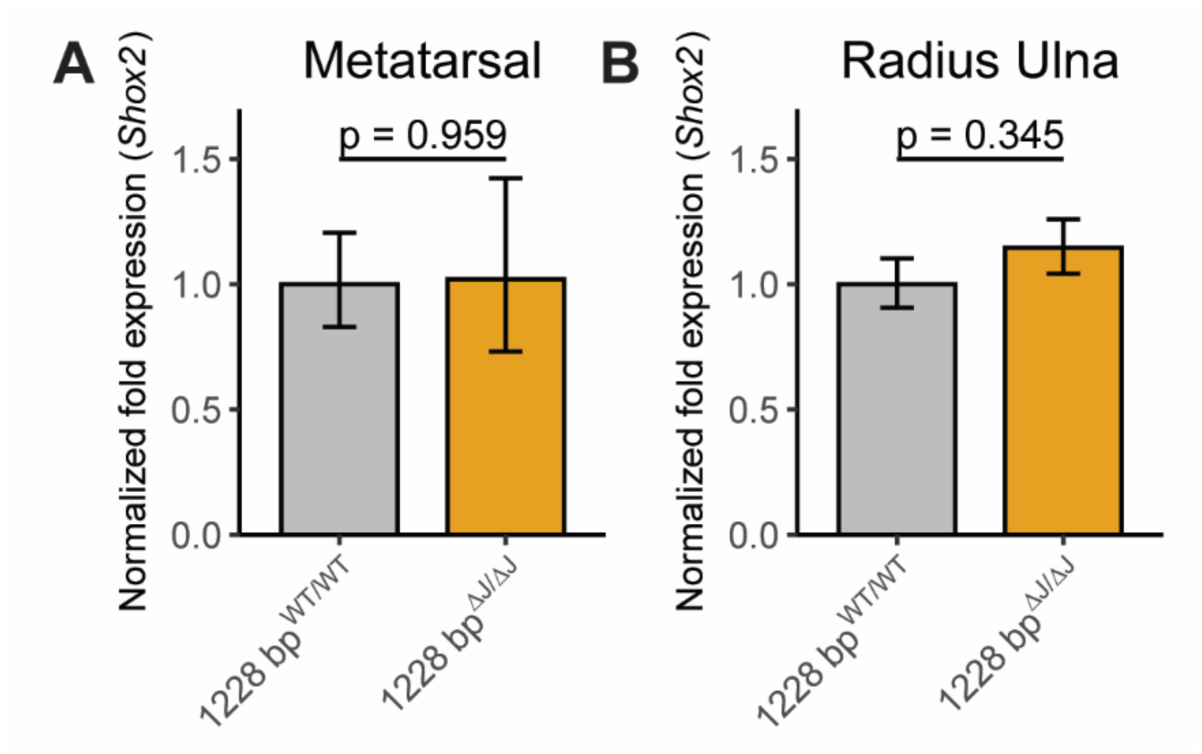
*Shox2* mRNA expression is unchanged in modified mice. *Shox2* expression was quantified by qPCR in growth cartilages of A) metatarsal and B) radius/ulna of P5 1228bp^ΔJ/ΔJ^ mice and wild type littermates. Expression was normalized to three reference genes: *Sdha, Tbp, Actin*. Error bars indicate SEM. Statistical significance was determined by Student’s t-test. n=6 (1228bp^WT/WT^ MT), n=6 (1228bp^ΔJ/ΔJ^ MT), n=5 (1228bp^WT/WT^ RU), n=7 (1228bp^ΔJ/ΔJ^ RU).

We suspected these data might expose limitations of using draft genome assemblies for ATAC-seq analyses to achieve highly selective goals. As part of the Vertebrate Genomes Project Phase 1, the *Jaculus jaculus* genome was sequenced again using PacBio Sequel II CLR, 10X Genomics, BioNano, and Hi-C v1 technologies and achieved a chromosome scale assembly (Formenti et al., 2026) (Supplemental Table S2). Using MACS3 and IDR, we re-evaluated our previous ATAC-seq data with this updated near-complete and error-free genome (mJacJac1.mat.Y.cur). Comparing the two analyses, we observed an increase in mapped reads, from 90-92% to 96-97% (Supplemental Table S3) with a high degree of reciprocal peak calling in the two assemblies (Supplemental Table S4). Among these, we again identify the *Shox2*-associated peak as reproducible in jerboa metatarsals and not in radius/ulna (Table 1). However, applying these same criteria in the revised assembly genome-wide, we find a total of 7,446 peaks (11.8%) are significant and reproducible in metatarsals and not in radius/ulna. Ongoing comparative genomics will evaluate the sequence evolution underlying these peaks.

**Table 1.**
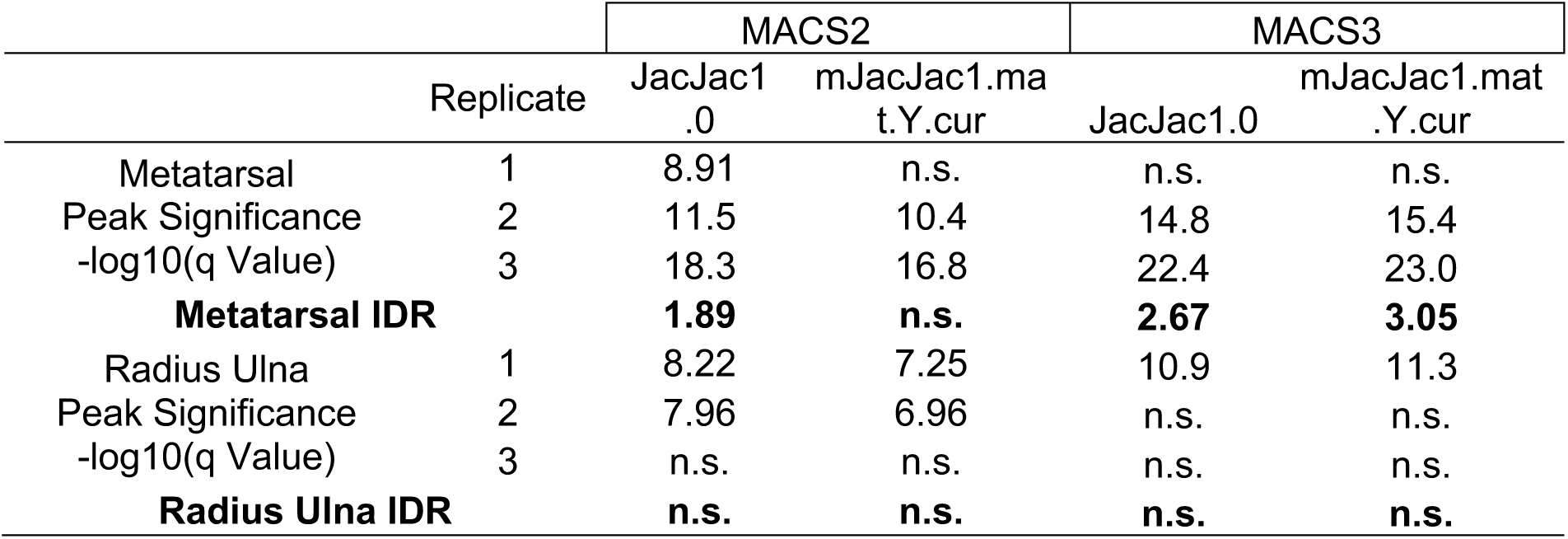
Comparison of ATAC-seq peak significance and IDR reproducibility scores (global IDR value) using the draft and revised jerboa genome assemblies. MACS2 used the mouse effective genome size as in Saxena et al, 2022 and MACS3 used calculated jerboa effective genome sizes; -log_10_(q value).

## Discussion

Genetic differences underlie the striking diversity of species forms and functions, but ‘how’ has been a mystery for more than a century. We now have an unprecedented number of genomes catalogued to uncover sequence differences, but we still cannot, in most cases, pinpoint the causative genetic basis of trait evolution.

In this study, we characterized the phenotypic effects on mouse limb elongation of introducing a single locus change identified in the lesser Egyptian jerboa, a rodent species with radically elongated hindlimbs. We previously identified an ATAC-seq peak that was accessible in jerboa metatarsals but not in radius ulna or in either cartilage of mice. The sequence underlying this peak is novel, resulting from the joining of sequence flanking a ∼1 kb deletion in the jerboa genome, and it is linked to a known skeletal growth regulator, *Short-stature homeobox 2* (*Shox2*). We further show that the ∼1 kb deletion is shared among bipedal jerboas and facultative bipedal jumping mice, and it removes sequence present in most of 488 other placental mammal genomes. This suggested that engineering the jerboa sequence difference in the mouse genome could be sufficient to alter limb proportion, but here we report that these mice have no detectable phenotype.

Considering the presumed genetic complexity of trait evolution, we did not expect that a single sequence modification would be sufficient to completely recapitulate jerboa metatarsal elongation in mice. The disproportionate growth that is responsible for diverse limb proportions undoubtedly results from many complex genetic interactions and not single-gene effects. Genome-wide association studies suggest that population variance in human height and body proportion is highly polygenic, or even ‘omnigenic’ (Chan et al., 2015; Lango Allen et al., 2010; Yengo et al., 2022), which presents a challenge to the classical genetics approach to identifying ‘mechanism’. Single locus changes have replicated macro-evolutionary traits, such as limb loss (Kvon et al., 2016) and tail loss (Xia et al., 2024), reinforcing a broader expectation of simple genetic cause. But trait loss can appear deceptively simple. Trait loss causes dedicated genetic networks to degenerate, and replicating the loss of function of a single gene that is *necessary* for trait development can appear *sufficient* to cause trait loss (Cooper, 2024).

Similar attempts to replicate constructive traits frequently resulted in subtle phenotypes, if any at all (Cretekos et al., 2008; Moreno et al., 2024; Ushiki et al., 2026). For example, replacing a single mouse limb enhancer that controls *Prx1*, a known endochondral growth regulator, with the orthologous sequence in the short-tailed fruit bat (*Carollia perspicillata*) caused a 6% forelimb elongation in neonates that lost statistical significance as inter-individual variance increased after birth (Cretekos et al., 2008). Similar results were observed using comparative genomics to understand the evolution of the patagium, a skin membrane extending between the limbs of gliding mammals (Moreno et al., 2024). Although *Emx2* expression is shown to be necessary (in sugar glider) and sufficient (in mouse) to recapitulate cell characteristics that are hallmarks of patagia outgrowth, none of three candidate glider accelerated enhancers changed *Emx2* expression or caused a phenotype when swapped into mice (Moreno et al., 2024).

The complex genetics of constructive traits further underscores the need for rigorous identification of candidate loci. Yet pinpointing the loci responsible for a given trait is inherently difficult, as the causative sequence changes are obscured by millions of years of neutral divergence or sequences under selection for different traits. This is the key challenge for every comparative genomic study.

Broadly speaking, two main approaches have narrowed the search space from a whole genome of billions of nucleotides to a tractable subset. The first relies on comparative sequence analyses to evaluate historically conserved noncoding elements (CNEs). High sequence conservation across a clade, as in across mammals, is frequently taken as evidence of functional importance (Christmas et al., 2023; Woolfe et al., 2005; Pennacchio et al., 2006). CNEs that are constrained across a clade but show a lineage-specific excess of substitutions are interpreted as candidates for recent functional divergence, as in human accelerated regions (HARs) (Pollard et al., 2006; Whalen & Pollard, 2022) and bat accelerated regions (BARs) (Booker et al., 2016). However, regulatory sequences can turn over substantially over evolutionary time without altering function (Phan et al., 2025; Kaplow et al., 2022; Villar et al., 2015), and conservation and/or selection at the sequence level is not, on its own, evidence of functional relevance to a particular trait. The second approach addresses both of these limitations. Rather than inferring function from sequence conservation, experimental methodologies (ATAC-seq, ChIP-seq, Hi-C, etc.) detect molecular signatures of regulatory activity directly and can expand our ability to identify potentially causative regulatory loci functioning in a tissue of interest and at the time of its divergent development (Gasperini et al., 2020; Hardison & Taylor, 2012). However, access to relevant tissues in non-traditional species is often a limitation.

Regardless of the approach to filter candidate regulatory sequences from the whole genome, whether using functional data or the set of evolutionarily conserved non-coding sequences, studies then identify potentially causative sequences by quantifying the relative rate of sequence change over evolutionary time. For example, a higher rate of nucleotide substitution relative to neutral drift or greater than the historical pattern of high conservation (accelerated) highlights sequences that may have experienced selection (Hubisz & Pollard, 2014). Sequence acceleration that correlates with evolution of a derived trait, especially in cases of convergent evolution, may be compelling causative candidates. However, as demonstrated here and in other studies (Roscito et al., 2021; Moreno et al., 2024), even sequences with strong evolutionary signatures and promising functional correlations may have no effect when singularly placed in another species.

Further, functional change may not only be caused by sequence evolution above the rate of drift. Even single nucleotide variants, which would not rise above the background rate of neutral substitution, can have enormous phenotypic consequences, as in the ZRS enhancer of *Shh.* A marginal increase (1.6-fold) in the transcription factor binding affinity of a low-affinity binding site is sufficient to cause polydactyly in humans and in mice (Lim et al., 2024). Although single nucleotide changes that have such a large phenotypic effect may be less likely to be the main drivers of evolution (Fisher, 1930; Cooper, 2024), they illustrate how subtle sequence changes that would not be detected by evolutionary rate comparisons do contribute substantially to phenotypic change while often major sequence changes, even to ultraconserved elements, do not (Ahituv et al., 2007; Snetkova et al., 2022).

The fragmented and incomplete state of the first jerboa genome precluded a comprehensive search of open chromatin regions and gene linkage, which limited our investigation to ‘cherry-picking’. We previously used ATAC-seq to identify putative functional *cis-* regulatory elements in limb growth cartilage and identified a substantial sequence change, a deletion, in the jerboa clade within one putative ATAC-seq peak. We chose to focus on this single peak because of its compelling narrative, but we detected no phenotype in mice that recapitulate the genetic difference. We cannot discern whether this sequence has no contribution to evolution of skeletal proportion or if it makes a small-effect contribution to a highly polygenic trait.

The genetic basis of trait evolution is likely extraordinarily complex, shaped by many genomic changes with small effects on phenotype that may not be detectable when modeled individually. Candidate-based and genome-wide comparative genomic approaches each offer valuable evolutionary insights, but each has a different shortcoming. Finding the evolutionary drivers of trait emergence therefore requires thoughtful integration of functional and comparative genomics and an understanding that our standard genotype-phenotype test of mechanism will frequently fail to replicate an evolved trait, regardless of the strength of evidence for the candidate sequence. However, though the definitive mechanisms underlying macroevolutionary change may remain out of reach, the effort will yield valuable insight into genome evolution and previously unknown gene functions.

## Acknowledgements

We are grateful to Dr. Aditya Saxena for early discussion of the sequence replacement strategy, to the laboratory of Dr. Emma Farley for assistance with the skeletal staining protocol, and to Dr. Daniel Richard for advice on the ATAC-seq analysis pipeline. We thank the UC San Diego Moores Cancer Center Transgenic and CRISPR Mouse Core for their assistance in generating modified mice. This work was supported by the National Institute of General Medical Sciences training grant (T32 GM133351) awarded to G.G. and by the National Institute of Arthritis, Skin, and Musculoskeletal Diseases under award number R01AR075415 and the National Science Foundation under award number IOS-1846390 to K.L.C. and by the Wu Tsai Human Performance Alliance and the Joe and Clara Tsai Foundation.

## Methods

All animal care and use protocols were reviewed and approved by the Institutional Animal Care and Use Committee (IACUC) of the University of California, San Diego.

### Generation of 1228bp^ΔJ/ΔJ^ mice

We modified 1228bp^ΔJ/ΔJ^ mice to recapitulate the sequence under the identified jerboa ATAC-seq peak, which consists of a ∼1 kb deletion and insertion of the 139 bp sequence under the peak. We performed this modification in C57BL/6 mice using two gRNAs cutting 151 bp upstream (chr3:67266338) and 56 bp downstream (chr3:67267772) from the peak located in an *Rsrc1* intron. The sequence associated with the jerboa ATAC-seq peak was inserted by homology-directed repair using a custom 560 bp ssODN with 90 bp homology arms that also recode the PAM sequences so that they cannot be cut again. We confirmed genotypes of founder mice by PCR and Sanger sequencing. Subsequent mice were genotyped by PCR.

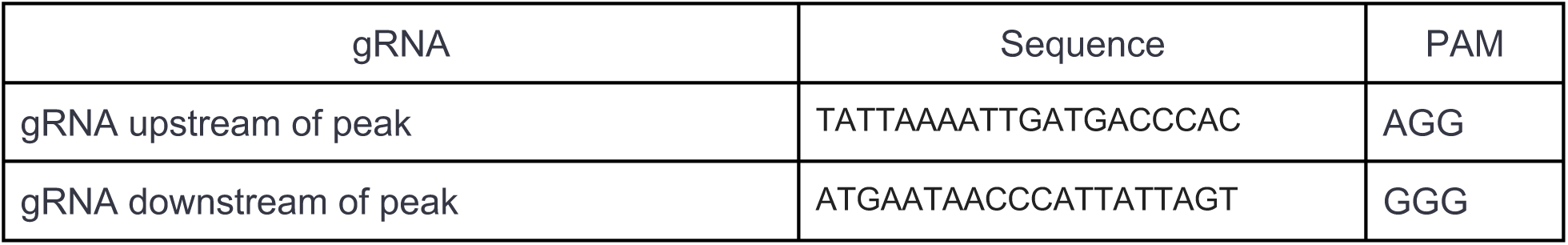

### Genotyping

Genomic DNA was extracted from tail or ear tissue and used to genotype modified mice with the following primers: Forward (5’-TCACTGGGGAGGCAGAAACG -3’), 1228bp^ΔJ^ Reverse (5’-CAGCAGCTGAATGCTTACATGG -3’), and WT Reverse (5’-TGACCACAAACAGCCTCCTC -3’). PCR was performed with an initial denaturation at 95°C for 3 min, followed by 30 cycles of denaturation at 95°C for 15 s, annealing at 58°C for 15 s, and extension at 72°C for 90 s, with a final extension at 72°C for 3 min. Expected band sizes are as follows:

| Primers | 1228bp<sup>ΔJ</sup> | 1228bp<sup>WT</sup> |
| --- | --- | --- |
| Forward + 1228bp<sup>ΔJ</sup> Reverse | ~ 1 kb | No band |
| Forward + WT Reverse | No band | ~ 500 bp |

### Micro-computed tomography (μCT) of limbs and skull and skeletal measurements

We collected 1228bp^ΔJ/ΔJ^ males and their wild type littermate controls at various ages (P14, P28, P42). The skulls, forelimbs, and hindlimbs were dissected, fixed, and stored in 70% ethanol at 4°C before scanning. We scanned animals by μCT at (18 μm)^3^ or (36 μm)^3^ isotropic voxel resolution (SkyScan 1076; SkyScan. Kontich, Belgium; 50 kVp, 200 μA, 0.5 mm aluminum filter, 180° scan, Δ=0.5°). We used plastic pellets to support specimens during scanning. After scanning, specimens were stored in 70% ethanol. Images are reconstructed using NRecon (Bruker, Belgium) with a smoothing factor of 1, ring artifact reduction factor of 6, beam hardening correction factor of 40%, and with a dynamic range from 0 to 0.11 attenuation units.

We measured the lengths of skeletal elements in the hindlimb (femur, tibia, metatarsals) and forelimb (humerus, radius, ulna, third metacarpal) from joint to joint, and skull (naso-occipital distance) using an academic license of the DragonFly 3D World software (Comet Technologies Canada Inc, 2025). We obtained all measurements blindly and present them as an average of left and right limb lengths.

### P0 and P7 skeletal stains and measurements

We collected 1228bp^ΔJ/ΔJ^ mice and wild type littermates at birth (P0) and post-natal day 7 (P7). Mice were skinned, eviscerated, and stored in 95% ethanol on a rocker overnight at room temperature. Skeletons were stained in cartilage staining solution (76% ethanol, 20% acetic acid, and 0.05% alcian blue 8GX) over two nights. Specimens were then incubated in 95% ethanol overnight, followed by 0.8% KOH overnight. Samples were subsequently incubated in bone staining solution (1% KOH, 0.005% alizarin red S) overnight. After staining, the skeletons were quickly rinsed with ddH2O and cleared in 1% KOH/20% glycerol over multiple nights until sufficiently cleared. Samples were then incubated 1% KOH/50% glycerol overnight, followed by 1% KOH/80% glycerol overnight, and finally stored in 100% glycerol. All overnight incubations were placed on a rocker at room temperature.

Following sufficient stain, we scanned skeletal preps on an Epson Perfection V600 Photo Scanner at a resolution of 1200 dpi. We measured lengths of the forelimb and hindlimb elements, from joint surface to joint surface, and the naso-occipital distance of each skull from digital scan images using FIJI ImageJ. Each skeletal element was blindly scanned and measured twice. Lengths were calculated as the average of each replicate scan, then averaged between left and right limbs.

### qPCR

We collected radius/ulna and metatarsal growth plates from P5 1228bp^ΔJ/ΔJ^ mice and 1228bp^WT/WT^ control littermates and stored them in RNAlater at 4°C overnight. RNAlater was removed the next day and growth plates were stored at -80°C. At the moment of RNA extraction, we froze growth plates in liquid nitrogen, disrupted them with mortar and pestle, followed by a further disruption with Qiagen Qiashredder. We then extracted RNA using the Qiagen RNAeasy Micro Kit. We generated cDNA using Invitrogen SuperScript III First-Strand cDNA synthesis kit following RNA normalization from simultaneously processed samples. We used 10 ng of cDNA to perform qPCR using SsoAdvanced™ Universal SYBR® Green Supermix. We measured *Shox2* expression levels using *Shox2* qPCR primers [*Shox2* Fwd (5’-3’): ACGGAGAGTGTCCCCTGAACT; and *Shox2* Rev, (5’-3’): CGCCTCTGCTTGATTTTGGT] (Cobb & Duboule, 2005) and normalized to *Actin* [*Actb* forward, TAATTTCTGAATGGCCCAGGTCT; and *Actb* Rev, ATTGGTCTCAAGTCAGTGTACAGG] (Bobick & Cobb, 2012), *Succinate Dehydrogenase Complex Flavoprotein Subunit A* [*Sdha* Fwd (5’-3’): GGAACACTCCAAAAACAGACCT; and *Sdha* Rev, (5’-3’): CCACCACTGGGTATTGAGTAG] (Saxena et al., 2022), and *TATA-box binding protein* [*Tbp* Fwd (5’-3’): CCGTGAATCTTGGCTGTAAACTTG, and *Tbp* Rev, (5’-3’): GTTGTCCGTGGCTCTCTTATTCTC](Bobick & Cobb, 2012). We performed all qPCR reactions under the same conditions (40 cycles, 60°C binding temp) in BioRad CFX Opus 96 Real-Time PCR System. We performed qPCR analysis in R following a previously described workflow (Taylor et al., 2019). Data is based on at least five animals per genotype (n=6 (1228bp^WT/WT^ MT), n=6 (1228bp^ΔJ/ΔJ^ MT), n=5 (1228bp^WT/WT^ RU), n=7 (1228bp^ΔJ/ΔJ^ RU)) and at least three technical replicates of each. In all replicates, 1228bp^ΔJ/ΔJ^ mice and 1228bp^WT/WT^ samples were simultaneously processed.

### Multiple sequence alignment

The multiple alignment of the jerboa ATAC-seq peak locus was created by fusing a 470-way mammal alignment and a newly created 73-way rodent alignment, which includes the revised *J. jaculus* assembly and additional jerboa species assemblies. First, the peak orthologous region was identified in the human hg38 assembly by liftOver. The sub-alignment including this region was extracted from the hg38-reference 470-way mammal alignment using UCSC Table browser. Then, the peak orthologous sequence was identified for each species based on the aligned genomic range in each species. Monotreme and marsupial assemblies were excluded because a large fraction of the peak locus does not align. The orthologous sequence for each rodent species was identified from the 73-way alignment following the same procedure. These orthologous sequences were re-aligned using Progressive Cactus (docker image: quay.io/comparative-genomics-toolkit/cactus, tag:v2.9.8) with default options. The resulting HAL alignment was converted to an mm10-reference MAF format by cactus-hal2maf using "--refGenome mm10 --chunkSize 10000 --dupeMode single --noAncestors”. The resulting MAF alignment was converted to fasta format and visualized along a phylogenetic tree using TreeViewer.

### ATAC-seq analysis

Two lanes per technical replicate were pooled as FASTQ files, and TrimGalore v0.6.10 was used to check read quality and to remove adapters. Jerboa reads were aligned to both JacJac1.0 and mJacJac1.mat.Y.cur genome assemblies with Bowtie2 v2.3.2 using local read alignment mode (--sensitive-local). The aligned reads were filtered for duplicates using Picard MarkDuplicates (docker image:broadinstitute/picard, tag:latest). BAM files of two technical replicates were pooled for each sample. BAM files were subsequently used for peak calling using MACS2 v2.1.1.2, with the following flags for ‘callpeak’: --format BAMPE --nolambda -- gsize mm as in (Saxena et al., 2022). The same BAM files were also used for peak calling with MACS3 v3.0.3, with the same flags except for --gsize: 2,470,267,327 for JacJac1.0 and 2,850,162,481 for mJacJac.mat.Y.cur. These values were each calculated by subtracting the number of ambiguous bases (Ns) from the total length of the genome assembly. Peaks reproducible across biological replicates were identified using IDR v2.0.4.2 with a global IDR value threshold of 0.05. Jerboa metatarsal peaks that do not overlap with any radius/ulna peaks are defined as metatarsal ‘unique’ peaks using bedtools v2.31.0 with ‘subtract -A’. To generate chains required for liftOver, a pair-wise whole-genome alignment of the two jerboa assemblies was produced using make_lastz_chain v3.1.7 (https://github.com/hillerlab/make_lastz_chains) with default parameters. liftOver was performed with --minMatch=0.1 -multiple -noSerial to allow for mapping to multiple loci. Two peaks that are independently identified in the two assemblies were defined as common if they are reciprocally mapped to each other by liftOver and overlap by at least 1 bp.

## Supplemental Figures

**Figure S1.**
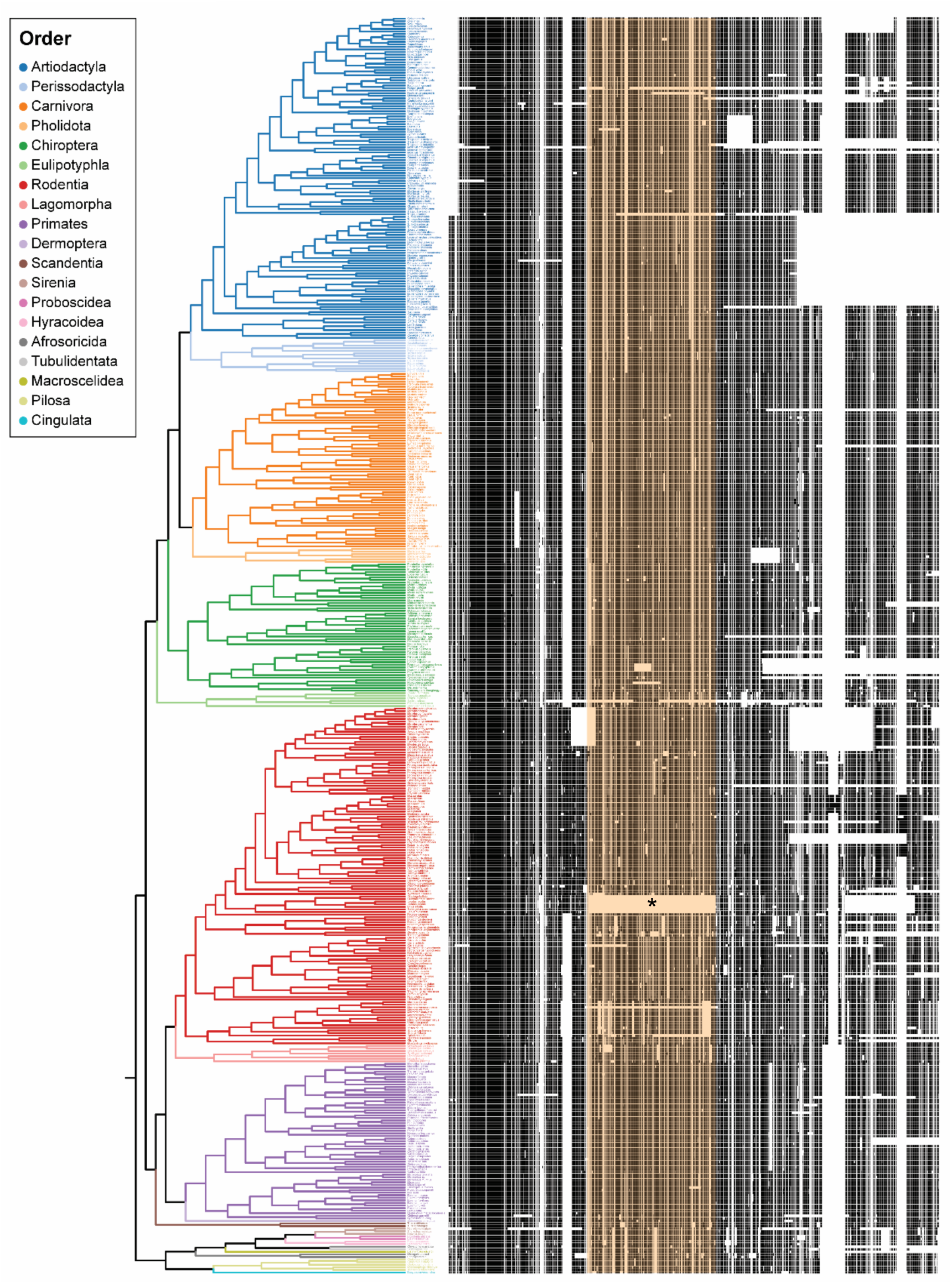
The ∼1 kb deletion is uniquely shared in the *Dipodoidea* superfamily. Alignment of sequence flanking the identified ATAC-seq peak in 488 mammals, including 131 rodent species, reveals a 1089 bp deletion (highlighted in orange) that is shared only within the *Dipodoidea* superfamily that includes jerboas and jumping mice (asterisk). Although other deletions within this sequence are present in a few mammal species, these are smaller and do not join the same flanking sequence.

**Figure S2.**
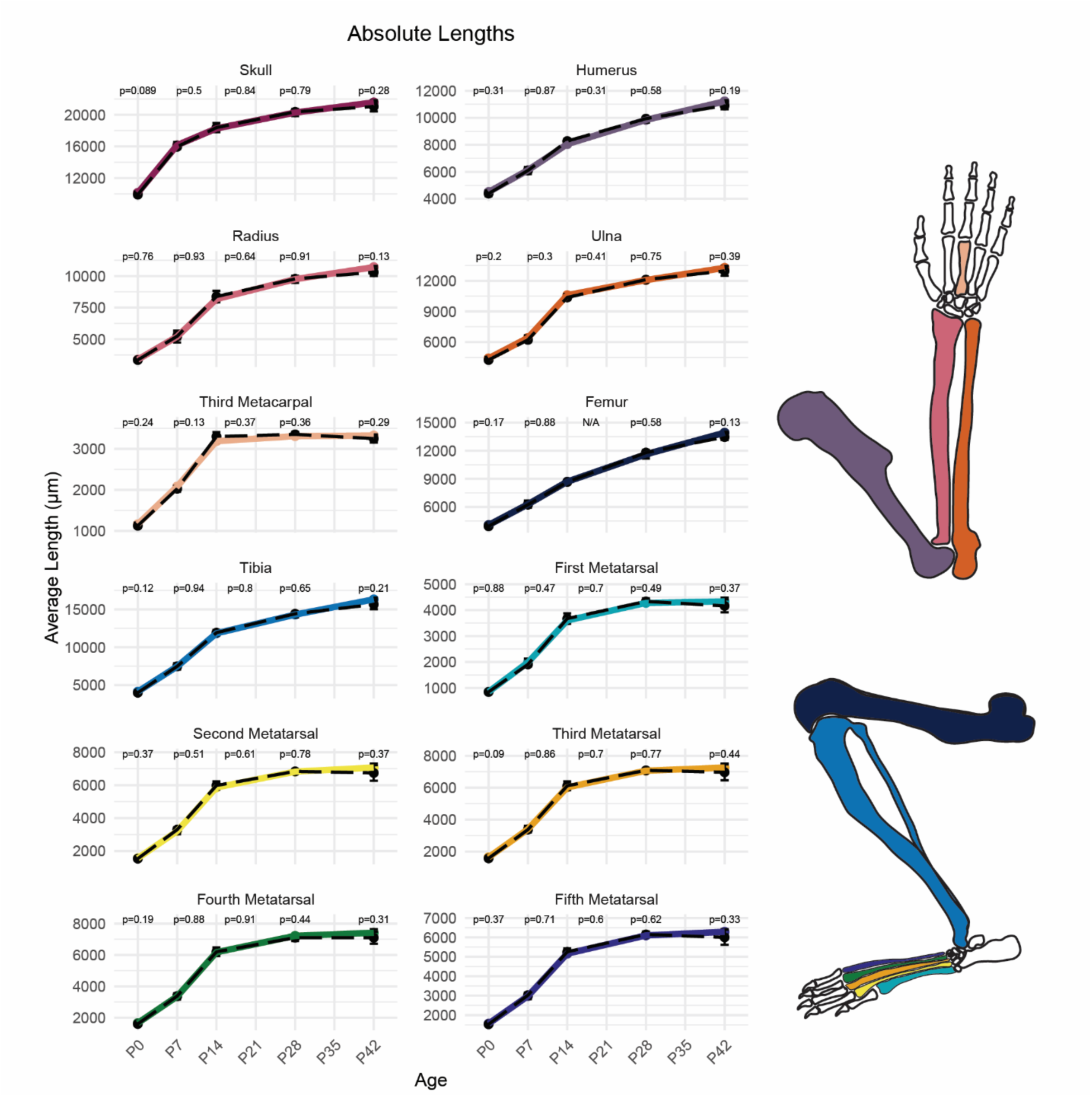
The absolute lengths of each limb bone show there is no detected limb skeletal phenotype. Lengths of the skull and limb skeletal elements were measured in modified 1228bp^ΔJ/ΔJ^ mice and wild type littermates at intervals from birth (P0) to adulthood (P42). Each measurement for one animal is the average length of the left and right limb skeletal elements. Wild type growth curves are represented in a black dashed line; 1228bp^ΔJ/ΔJ^ growth curves are presented as a solid color line. Statistical significance was determined by Student’s t-test. n≥ 3

**Figure S3.**
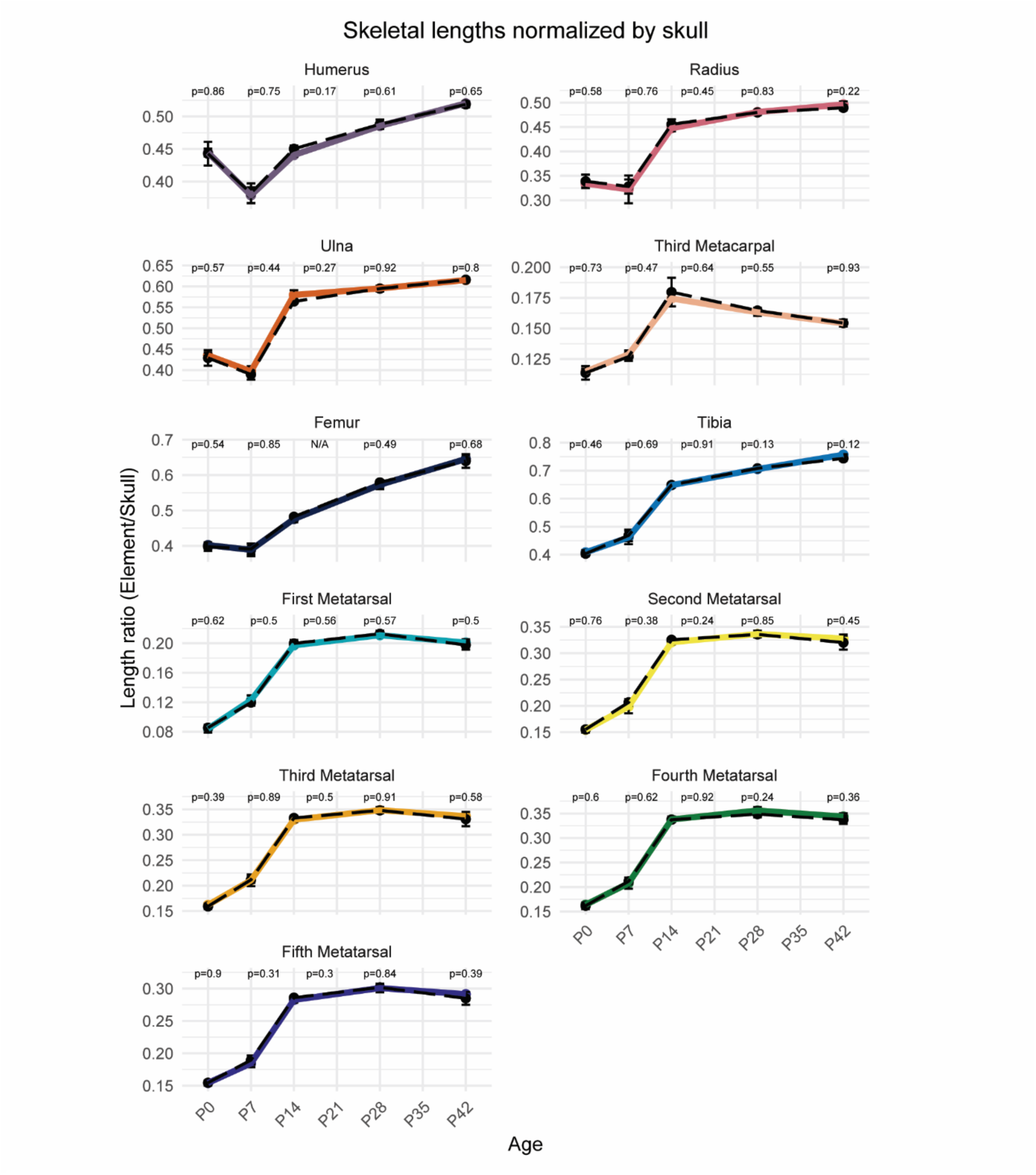
The lengths of each limb bone normalized to the naso-occipital length of the skull reinforces there is no detectable limb phenotype. Lengths of the skull and limb skeletal elements, normalized to skull length, in modified 1228bp^ΔJ/ΔJ^ mice and wild type littermates at intervals from birth (P0) to adulthood (P42). Each measurement for one animal is the average length of the left and right limb skeletal elements divided by the length of the skull for the same animal. The observed decreased normalized ratio at P7 is due to disproportionally accelerated elongation of the skull at this timepoint (see Figure S2). Wild type growth curves are represented in a black dashed line; 1228bp^ΔJ/ΔJ^ growth curves are presented as a solid color line. Statistical significance was determined by Student’s t-test. n≥ 3 Genotype ratios resulting from 1228bp^ΔJ/WT^ x 1228bp^ΔJ/WT^ crosses

**Table S1.** Intercrosses of 1228bp^ΔJ/WT^ mice produce genotype frequencies that that are not significantly different from expected by Mendelian inheritance.

| Genotype ratios resulting from 1228bp<sup>ΔJ/WT</sup> x 1228bp<sup>ΔJ/WT</sup> crosses |  |  |  |
| --- | --- | --- | --- |
| Genotype | 1228bp<sup>WT/WT</sup> | 1228bp<sup>ΔJ/WT</sup> | 1228bp<sup>ΔJ/ΔJ</sup> |
| Number of pups | 89 | 140 | 79 |
| Ratio | 0.289 | 0.455 | 0.256 |

**Table S2.** Comparison of assembly details for draft and revised jerboa genome assemblies used for ATAC-seq.

|  | JacJac1.0 | mJacJac1.mat.Y.cur |
| --- | --- | --- |
| Year | 2012 | 2021 |
| Genome size | 2.8 Gb | 2.9 Gb |
| Total ungapped length | 2.5 Gb | 2.9 Gb |
| Number of chromosomes | N/A | 25 |
| Number of scaffolds | 10,899 | 161 |
| Scaffold N50 | 22.1 Mb | 158.2 Mb |
| Scaffold L50 | 36 | 7 |
| Number of contigs | 337,361 | 715 |
| Contig N50 | 15.7 kb | 22.1 Mb |
| Contig L50 | 44,339 | 39 |
| GC percent | 42 | 42 |
| Genome coverage | 78x | 61x |
| Assembly level | Scaffold | Chromosome |
| Ambiguous bases ('Ns') | 364,982,898<br>(12.9%) | 13,703,780<br>(0.478%) |
| Sequencing technology | Illumina Hi-Seq | PacBio Sequel II<br>Continuous Long<br>Read |

**Table S3.** Comparison of ATAC-seq read alignment and significant peaks detected using the draft and revised jerboa genome assemblies.

| Sample | Replicate | % ATAC reads aligned |  | Number of ATAC peaks |  |
| --- | --- | --- | --- | --- | --- |
|  |  | JacJac1.0 | mJacJac1.mat.Y.cur | JacJac1.0 | mJacJac1.mat.Y.cur |
| Metatarsal | 1 | 90.8% | 97.4% | 172,857 | 165,409 |
|  | 2 | 92.7% | 97.8% | 168,492 | 194,698 |
|  | 3 | 92.2% | 97.7% | 168,251 | 199,773 |
| Radius<br>Ulna | 1 | 92.0% | 97.8% | 219,887 | 206,696 |
|  | 2 | 92.3% | 97.6% | 192,201 | 221,086 |
|  | 3 | 90.9% | 96.7% | 202,815 | 191,278 |

**Table S4.** Comparison of peak sequence alignment and peak overlap using the draft and revised jerboa genome assemblies.

|  | JacJac1.0 | mJacJac1.mat.Y.cur |
| --- | --- | --- |
| Total number of IDR peaks | 67,842 | 63,236 |
| % with >10% sequence alignment in reciprocal genome | 97.9 | 98.3 |
| % of peaks also called in reciprocal genome | 84.8 | 88.3 |

## Notes

### Competing Interest Statement

The authors have declared no competing interest.

